# Sex-specific actions of estradiol and testosterone on human fibroblast and endothelial cell proliferation, bioenergetics, and vasculogenesis

**DOI:** 10.1101/2023.07.23.550236

**Authors:** Ashley T. Martier, Yasmin V. Maurice, K. Michael Conrad, Franck Mauvais-Jarvis, Mark J. Mondrinos

**Affiliations:** Department of Biomedical Engineering, Tulane University School of Science & Engineering, New Orleans, LA, USA; Tulane Center for Excellence in Sex-based Biology and Medicine, New Orleans, LA, USA; Section of Endocrinology, Deming Department of Medicine, Tulane University School of Medicine, New Orleans, LA, USA; Southeast Louisiana VA Medical Center, New Orleans, LA, USA; Department of Physiology, Tulane University School of Medicine, New Orleans, LA, USA

## Abstract

Progress toward the development of sex-specific tissue engineered systems has been hampered by the lack of research efforts to define the effects of sex-specific hormone concentrations on relevant human cell types. Here, we investigated the effects of defined concentrations of estradiol (E2) and dihydrotestosterone (DHT) on primary human dermal and lung fibroblasts (HDF and HLF), and human umbilical vein endothelial cells (HUVEC) from female (XX) and male (XY) donors in both 2D expansion cultures and 3D stromal vascular tissues. Sex-matched E2 and DHT stimulation in 2D expansion cultures significantly increased the proliferation index, mitochondrial membrane potential, and the expression of genes associated with bioenergetics (Na+/K+ ATPase, somatic cytochrome C) and beneficial stress responses (chaperonin) in all cell types tested. Notably, cross sex hormone stimulation, i.e., DHT treatment of XX cells in the absence of E2 and E2 stimulation of XY cells in the absence of DHT, decreased bioenergetic capacity and inhibited cell proliferation. We used a microengineered 3D vasculogenesis assay to assess hormone effects on tissue scale morphogenesis. E2 increased metrics of vascular network complexity compared to vehicle in XX tissues. Conversely, and in line with results from 2D expansion cultures, E2 potently inhibited vasculogenesis compared to vehicle in XY tissues. DHT did not significantly alter vasculogenesis in XX or XY tissues but increased the number of non-participating endothelial cells in both sexes. This study establishes a scientific rationale and adaptable methods for using sex hormone stimulation to develop sex-specific culture systems.

## Introduction

Biological sex is defined by the differences between males and females in sex chromosomes, sex organs, and sex hormones. Sex chromosome composition, sex hormone concentrations, and epigenetic changes induced by the fetal testosterone surge in male cells all contribute to sex-specific differences in cell biology, tissue and organ physiology, pathophysiology, and response to treatments (1). While conditions such as Kleinfelter syndrome demonstrate that biological sex is not binary (2), we refer to female cells as containing XX chromosomes and male cells as containing XY chromosomes and we use the term sex to refer to these differences in the context of our in vitro model development. Gender is distinct from sex and their convergence modifies health and disease, but it is more difficult to isolate the effect of gender in preclinical research (3). Biomedical research has historically overlooked sex as a critical factor in study design, and the National Institutes of Health (NIH) has mandated the inclusion of sex as a biological variable (SABV) in all NIH-funded studies (4). This shift in policy is significant as a majority of publications in fields such as tissue engineering and microphysiological systems (MPS) have historically neglected to report the sex of cells used or consider sex-specific study design (5).

Sex-specific tissue and organ models are urgently needed to help decipher mechanisms underlying sex differences observed in pathologies such as cancers, cardiovascular diseases, and organ fibrosis (6–8). The relative contributions of genetic differences and environmental cues, such as hormone levels, to the observed sexual dimorphisms remains unclear (9). The sex of cells influences genetic profiles across the genome, suggesting that there are innate genetic and epigenetic differences between male and female cells beyond the obvious differences in sex chromosome composition (10). Clinical observations, such as the increased incidence of cardiovascular diseases in post-menopausal females with decreased estrogen levels, hints that hormones play a vital role in observed sexual dimorphisms (11). Therefore, rational design of biological female and male tissue models will benefit from a more complete understanding of sex-specific hormone actions on cellular physiology in culture and how these effects influence morphogenesis and function in engineered models.

Biomimicry of the microvasculature has remained a critical pursuit in tissue engineering (12). Recently, engineering vascularized organ chips and microphysiological systems has unlocked deeper levels of physiological relevance for preclinical modeling (13–17). There is an established framework for manipulating biochemical and biophysical cues to guide the processes of vasculogenesis and angiogenesis in vitro, but none of these methods account for biological sex (18). There is a rationale to suspect that female and male vascular cells will respond differently to engineered culture environments. Expansion culture of human endothelial cells and vascular support cells such as fibroblasts, pericytes, and smooth muscle cells as primary cells or stem cell derivatives is a core biotechnology process in tissue engineering, yet sex-specific culture methods do not exist (19–23).

Here, we report sex-specific effects of E2 and DHT in defined media formulations for 2D expansion culture of primary human fibroblasts and endothelial cells and in a 3D bulk tissue vasculogenesis assay. Our data reveals that sex-matched hormone stimulation increases proliferation and bioenergetic capacity, whereas cross sex hormone stimulation exerts inhibitory effects. E2 treatment enhanced vasculogenesis in female tissues and inhibited vasculogenesis in male tissues, following 2D cultures. Conversely, DHT did not enhance vasculogenesis in male tissues and increased the number of non-interconnected endothelial cells in both female and male tissues. Collectively, our data informs the development of sex-specific primary cell culture systems and tissue engineering methods.

## Methods

### 2D Cell Culture

Primary human lung fibroblasts (HLF), human dermal fibroblasts (HDF), and human umbilical vein endothelial cells (HUVEC) from both XX and XY donors were sourced for use in these studies (ages 20-45, ATCC). HLF and HDF were expanded in fibroblast basal medium supplemented with fibroblast growth kit – low serum (ATCC) and HUVEC were grown in vascular cell basal medium supplemented with endothelial growth kit – VEGF (ATCC). At 50% confluence, cells were changed to phenol red-free media supplemented with charcoal stripped serum (Gibco) for a 24 hour serum starvation. After 48 hours of starvation, we exposed cells to 0nM, 1nM or 10nM E2 dissolved in ethanol (Thermo Fisher). E2 was replenished every 24 hours and media was changed every 48 hours. At day 1, 2, 3, 4, and 7, cells were collected via trypsinization. These trials were repeated with 0nM, 1nM, and 10nM DHT using the same time points and methodology as was used for E2 exposures. Cell pellets were stored in RNAlater (Thermo Fisher) at 4°C until RNA isolation.

### Gene expression analysis

mRNA was isolated from collected samples via the Qiagen RNeasy kit following the supplier protocol, and sample purity was validated by measuring the ratio of absorbance at 260 and 280 nm (260/280) Nanodrop spectrophotometry (Thermo Fisher). mRNA samples were stored at −80°C for long term storage. cDNA was synthesized using qScript cDNA SuperMix (QuantaBio) according to the supplier recommended protocol, diluted 1:10 in UltraPure DNase and RNase free water (Thermo Fisher), and stored at −20°C. Quantitative reverse transcription polymerase chain reaction (qRT-PCR) was used to measure the expression of genes associated with energetics and stress response in all cells. qRT-PCR was completed using SybrGreen PowerUp on the ThermoFisher StepOnePlus system (Thermo Fisher). Primers used are listed in **Table 1**. GAPDH was utilized as the housekeeping gene. PCR results were analyzed via the 2^-ΔΔCt^ method.

### Fabrication of PDMS microarrays and surface modification for hormone culture

A 9-well milliscale tissue array was designed in SolidWorks (Dassault Systèmes) and transferred into PreForm 3D Printing Software (FormLabs). Molds were printed using a Form3 SLA 3D printer (FormLabs). As reported elsewhere, after printing the molds were washed, cured, and flattened to generate acceptable molds for PDMS soft lithography (ACS paper). As PDMS is known to absorb small, hydrophobic molecules such as steroid hormones (24), molds were polyurethane coated to increase smoothness of PDMS and reduce surface area to minimize absorption. Absorption of sex hormones was further mitigated by cross-mixing PDMS-PEG self-segregating polymers into the PDMS mixture prior to PDMS curing (25). The ideal amount of PDMS for our purposes was found to be 0.25% by weight as this endowed the material with the surface properties we desired while maintaining the benefits of traditional PDMS. As such, PDMS was mixed at a 1:10 ratio of elastomer to fixative agent (Sylgard 184, Ellsworth Adhesives). The PDMS-PEG copolymer was then added to a total of 0.25% weight (Gelest, DBE-712). The polymer was then mixed thoroughly and degassed. PDMS was poured into molds, degassed again, and kept in an oven at 60°C for at least 4 hours to ensure complete curing. Prior to use, we sanitized devices via UV light sterilization for 2 hours. To ensure complete attachment of the collagen gels to the PDMS devices, the wells of the arrays were treated with a 5 mg/ml polydopamine solution for 2 hours in UV light (26). Devices were then stored in a dark space at room temperature until use.

To test that PDMS-PEG copolymer microarrays would prevent hormone absorption, Nile Red absorption assays were completed. Nile Red, a fluorescent stain with similar size to E2, has been used in previous studies to assay PDMS absorption of small molecules (24). 1μg/mL Nile Red (Thermo Fisher) was dissolved in ethanol and loaded into central wells of small PDMS copolymer devices and allowed to sit for 1 hour. Devices were washed with PBS 3 times. PDMS with or without copolymer made on coated and non-coated molds were imaged under fluorescence. Images were quantified using the average gray intensity across the well as calculated in FiJi.

### 3-D vasculogenesis assay

HUVEC and HLF were expanded using the same techniques used in 2D culture. PDA treated PDMS microarrays were loaded with 2 x10^6^ cells/ml HLF and 2 x10^6^ cells/ml HUVEC in a hydrogel comprised of blended collagen type I (2.25 mg/ml collagen type I, Corning) and fibrin (5mg/mL fibrinogen activated with 1 U/ml thrombin, Sigma Aldrich). Hydrogels were allowed to solidify at 37°C for 30 minutes. Stromal vascular tissues were cultured in vascular cell growth media supplemented with VEGF (ATCC), 25 μg/mL aprotinin (Sigma Aldrich), and charcoal stripped serum (Gibco) for 48 hours. Media was changed to phenol red free media containing 2% charcoal stripped serum for complete hormone starvation for 24 hours before exposing the gels to 0nm, 1nm, or 10nm E2 in ethanol or 0nM, 1nM, or 10nM DHT for 72 hours.

### TMRM dye assay

The tetramethylrhodamine methyl ester (TMRM) dye was used as an indirect indicator of mitochondrial membrane potential and ATP generation capacity (27). To prepare monolayers, XX and XY HUVEC and HLF were seeded at a density of 10,000 cells per well in 24-well tissue culture plates and grown in expansion medium for 48 hours. After 48 hours, cells were switched to phenol red-free media supplemented with 2% charcoal stripped serum for 24 hours to ensure complete hormone starvation. Following starvation, cells were exposed to media supplemented with 1nM or 10nM E2 or 1nM or 10nM DHT dissolved in absolute ethanol. Control samples were exposed to media containing 0.1% ethanol as a vehicle control. For each condition, 3 wells were grown for each XX and XY cell line for n=3. Cultures were treated with hormones for 48 hours prior to exposing the cells to 100nM TMRM dissolved in DMSO for 30 minutes at 37°C as per manufacturer instructions (Thermo Fisher). Monolayers were washed with PBS and imaged immediately.

### Immunofluorescence and 3D whole mount staining

Monolayers of HUVEC and HLF were prepared as described above and then fixed with 4% paraformaldehyde at room temperature for 15 minutes. Monolayers were then gently washed with PBS 3 times. Fixed well plates were stored covered in PBS at 4°C until staining. Samples were blocked and permeabilized with 1% bovine serum albumin (BSA) and 0.1% Triton-X (Sigma Aldrich) in PBS at room temperature for 20 minutes. Rabbit anti-Ki67 antibody (Abcam) was added at 10 μL/mL in 0.1% BSA in PBS. Plates were rocked for 2 hours protected from light at a gentle rock and then washed with PBS 4 times. Secondary Alexa fluor 488 at 4μL/mL, 4 μL/mL 4′,6-diamidino-2-phenylindole (DAPI) to label nuclei, and 4 μL/mL phalloidins (Thermo Fisher) to label actin were added in 0.1% BSA in PBS for 40 minutes in a dark, gentle rocking. Plates were then washed again in PBS and stored covered in PBS at 4°C protected from light until imaging.

3D tissues from the PDMS tissue arrays were processed for whole mount staining. After 72 hours, tissues were fixed in 4% paraformaldehyde (Sigma Aldrich) for 1 hour at room temperature and then overnight at 4°C. Tissues were then washed with several changes of PBS and stored in PBS at 4°C until stained. Gels were stained using 4 μL/mL 4′,6-diamidino-2-phenylindole (DAPI) to label nuclei, 4 μL/mL phalloidins to label actin, and 20 μL/mL Ulex Europeas agglutinin I (UEA-1) to specifically label endothelial cell lectins. Stains were prepared in PBS with 0.2% Triton-X and 1% BSA (Sigma Aldrich). Devices were loaded with staining cocktail, ensuring complete coverage of the gels, and rocked gently for 1 hour at room temperature. Gels were then refrigerated overnight, rocked for an additional hour the next day at room temperature, washed in PBS, and stored covered with PBS at 4°C until imaging.

### Imaging and Image Analysis

All 2D and 3D samples were imaged on an inverted Nikon C2 laser scanning confocal microscope (LCSM) equipped with a Nikon DS-FI3 camera. Samples from a given staining cohort were imaged at a fixed laser intensity and exposure time. For 2D cultures, 5 images of each sample were taken to create 5 technical replicates. Analysis of 2D stains was performed in FiJi. TMRM images were converted to grayscale and segmented to the area of the cells utilizing an overlayed DIC image of the sample. Mean fluorescence intensity of the region of the cells was measured, and the average intensity across the image was recorded. This process was completed for each of the 5 images taken from each well. The average of these 5 measurements was then used as the average intensity across the entire well. Ki67 staining was analyzed in FiJi, using the DAPI co-stain as the mask for regions of interest (cell nuclei). The threshold intensity of DAPI images was adjusted to the sharply capture cell nuclei and turned into a binary mask which was overlaid on the Ki67 channel. Ki67 positive cells were counted via particle analysis.

Each tissue from the vasculogenesis assay was imaged at a 9×9 image array of full thickness Z-stacks to image the entire three-dimensional bulk volume of the tissues, excluding a rim of tissue at the interface with PDMS surfaces. 3D image datasets were analyzed via MATLAB (Mathworks, R2021b). Max intensity projections of the Z-stacks were saved at TIFF files and exported to MATLAB for filtering and quantification. Briefly, vascular images were smoothed with a gaussian filter, low intensity noise was filtered out, and images were then denoised via a pretrained neural network. The open-source segmentation tool REAVER was utilized to segment vascular networks and quantify morphometric parameters (28). Image analysis results were exported to GraphPad Prism V 9.2 for statistical analysis.

### Statistical Analysis

For E2 exposure PCR, HLF were repeated n=4 times, and HDF and HUVEC were repeated n=3 times. DHT exposure was repeated n=3 times. Statistical analyses on PCR results were completed via one-way ANOVA with a 95% confidence interval for significance (* = p < 0.05; ** = p < 0.01; *** = p < 0.001). For analysis of TMRM and Ki67 indexes, statistical analysis was completed via unpaired, two-tailed t-tests with a 95% confidence interval (* = p < 0.05; ** = p < 0.01; *** = p < 0.001). Details on sample sizes and numbers of replicates are included in previous methods sections and figure captions.

## Results

### Sex-Specific Hormone Effects on Fibroblast and Endothelial Cell Proliferation

We treated female (XX) and male (XY) human lung fibroblasts (HLF) and human umbilical vein endothelial cells (HUVEC) with estradiol (E2,10 nM) or dihydrotestosterone (DHT, 10 nM) for 48 hours and measured cell proliferation by staining for the cell cycle marker Ki67 (**Figure 1A, B**). The percentage of cells expressing Ki67 (Ki67 index) is commonly used to quantify cell proliferation (29, 30). The baseline Ki67 index of vehicle-treated HLF cultured in FGM expansion medium made with charcoal-stripped serum was 59.73+/-9.06% for XY and 65.95 +/-3.75% for XX (**Figure 1C**). DHT treatment significantly increased the Ki67 index to 92.50+/-5.66% in XY HLF (p<0.001). In contrast DHT produced no significant effect in XX HLF (**Figure 1C**). E2 treatment of XX HLF increased the Ki67 index to 92.52+/-2.11%, a significant increase of over 40% (p<0.01). In contrast, E2 produced no significant effect in XY HLF cells. In HUVEC, baseline Ki67 indexes were 70.81 +/- 6.97% for XX and 74.59+/- 4.71% for XY cells (**Figure 1D**). DHT treatment increased XY HUVEC proliferation with an increased Ki67 index of 86.49+/-1.86% (p<0.05) and decreased XX HUVEC proliferation to a Ki67 index of 57.33+/- 6.88% (p < 0.05). In contrast, E2 potently inhibited XY HUVEC proliferation with a decreased Ki67 index of 44.06 +/- 3.07% (p< 0.001) and increased XX HUVEC proliferation with a Ki67 index of 89.84+/-3.57% (p < 0.01). We also noted that E2 treatment qualitatively decreased the formation of organized actin cytoskeletal fibers in XY HUVEC and HLF (**Figure 1E**). Thus, in HLF and HUVEC, DHT and E2 exhibit sex-specific proliferative and antiproliferative effects.

**Figure 1.**
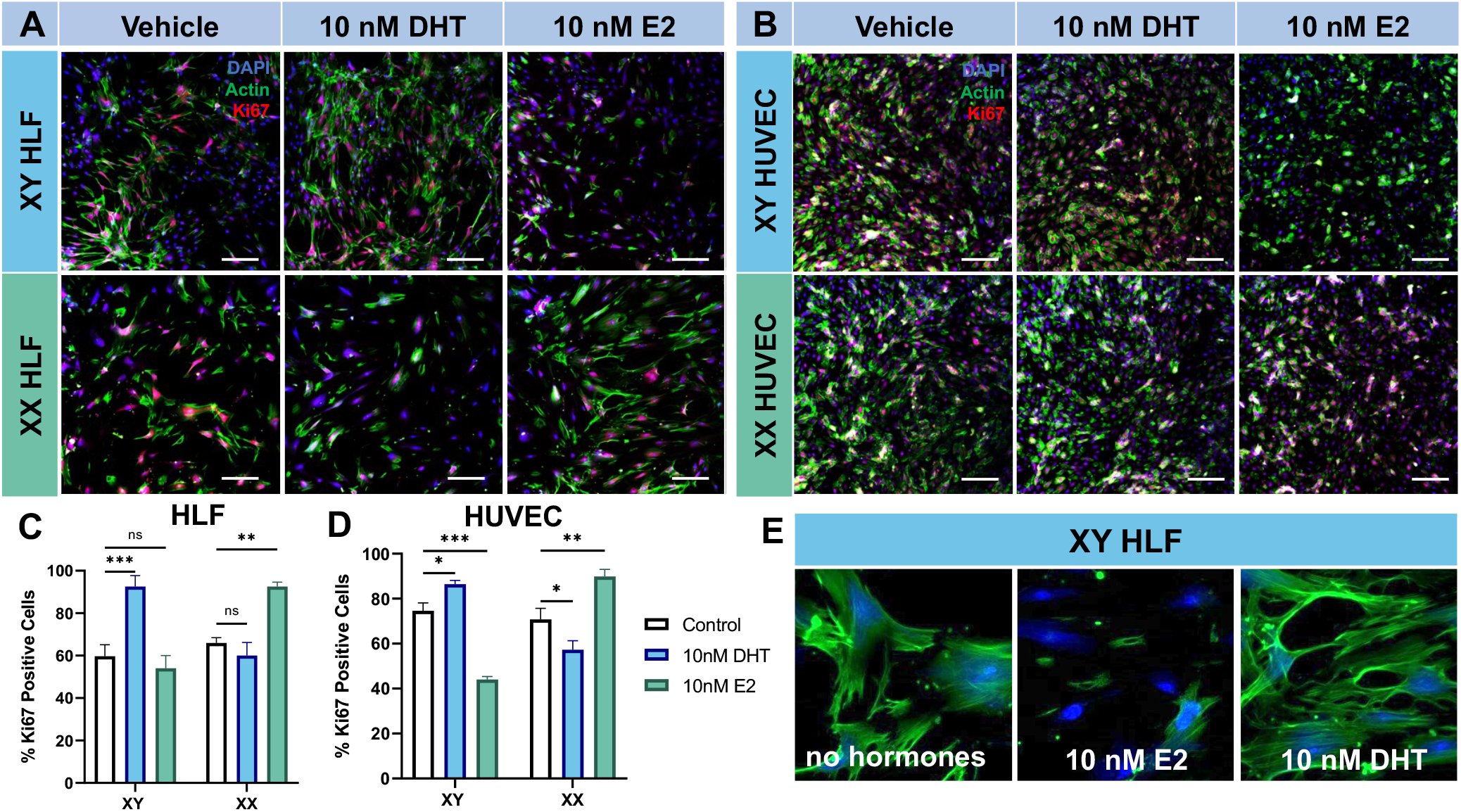
Sex-specific effects of E2 and DHT on endothelial cell and fibroblast proliferation. **A:** Representative micrographs of XY and XX HLFs exposed to 10nM DHT and 10nM E2 for 48 hours stained with DAPI (blue), Actin (green), and Ki67 (red). **B:** Representative micrographs of XY and XX HUVECs exposed to 10nM DHT and 10nM E2 stained for 48 hours with DAPI (blue), Actin (green), and Ki67 (red). Scale bar = 200 µm. **C:** XY HLFs demonstrated a significant increase in Ki67 positivity when exposed to 10nM DHT and a non-significant decrease in Ki67 expression when exposed to 10nM E2. XX HLFs demonstrated a non-significant decrease in Ki67 expression after 10nM DHT exposure and a significant upregulation in Ki67 positivity following exposure to 10nM E2. **D:** HUVECs reacted to hormone exposure in a similar way to HLFs, but XY cells experienced a significant decrease in signal after 10nM E2 exposure and XX cells experienced a significant decrease in positivity after 10nM DHT exposure. HLF and HUVEC levels compared against vehicle control groups via ANOVA (n=3 images per monolayer, 2 monolayers per condition). * = p < 0.05; ** = p < 0.01; *** = p < 0.001. **E:** Exposure of XY HLF to 10nM E2 caused disruption of normal actin fiber alignment.

### Estradiol Induces Sex-specific Alterations of Cellular Bioenergetics in 2D Culture

Cell proliferation increases energy demand and requires increased ATP production (31). Higher rates of proliferation also increases protein translation demands, potentially inducing endoplasmic reticulum stress unless the protein folding and trafficking machinery is upregulated (32). We sought to determine if sex-specific hormone concentrations (E2 stimulation of XX cells), which significantly increased cell proliferation (**Figure 1**), would upregulate the expression of key genes associated with ATP production and proteostasis, specifically the Na+/K+ ATPase (ATP1A1) that enables cells to utilize ATP for ion pumping, somatic cytochrome C (CYCS) that contributes to ATP generation via oxidative phosphorylation, and the chaperonin heat shock protein-60 (HSPD1) that aids in the management of increased translational burden. We also assessed if cross sex hormone stimulation (E2 stimulation of XY cells), which significantly decreased cell proliferation (**Figure 1**), would downregulate expression of ATP1A1, CYCS, and HSPD1. We chose this set of genes as a molecular readout for comparing E2 and DHT responses in multiple primary human cell types.

E2 significantly upregulated ATP1A1, CYCS, and HSPD1 in XX HLF cells by 24 hours; at this time point, both 1nM E2 and 10nM E2 elicited a significant increase in ATP1Acr1 while only 10nM E2 caused significant upregulation in CYCS and HSPD1 at 24 hours (**Figure 2A**). This upregulation was maintained significantly above baseline after 72 hours at both 1nM and 10nM E2 concentrations, after which expression levels dropped to near baseline levels. Of note, HSPD1 levels remained upregulated through 96 hours at both concentrations tested before returning to baseline. E2 treatment produced the same pattern of upregulation through 72 hours in XX HDF with a trend of returning to baseline, and 10nM E2 caused a greater upregulation in all 3 genes for a longer time frame than 1nM E2. Of note, in HDF, expression levels of ATP1A1 and CYCS were significantly elevated above baseline up until the day 7 endpoint, suggesting potential organ-specific effects of E2 stimulation (**Figure 2B**).

**Figure 2.**
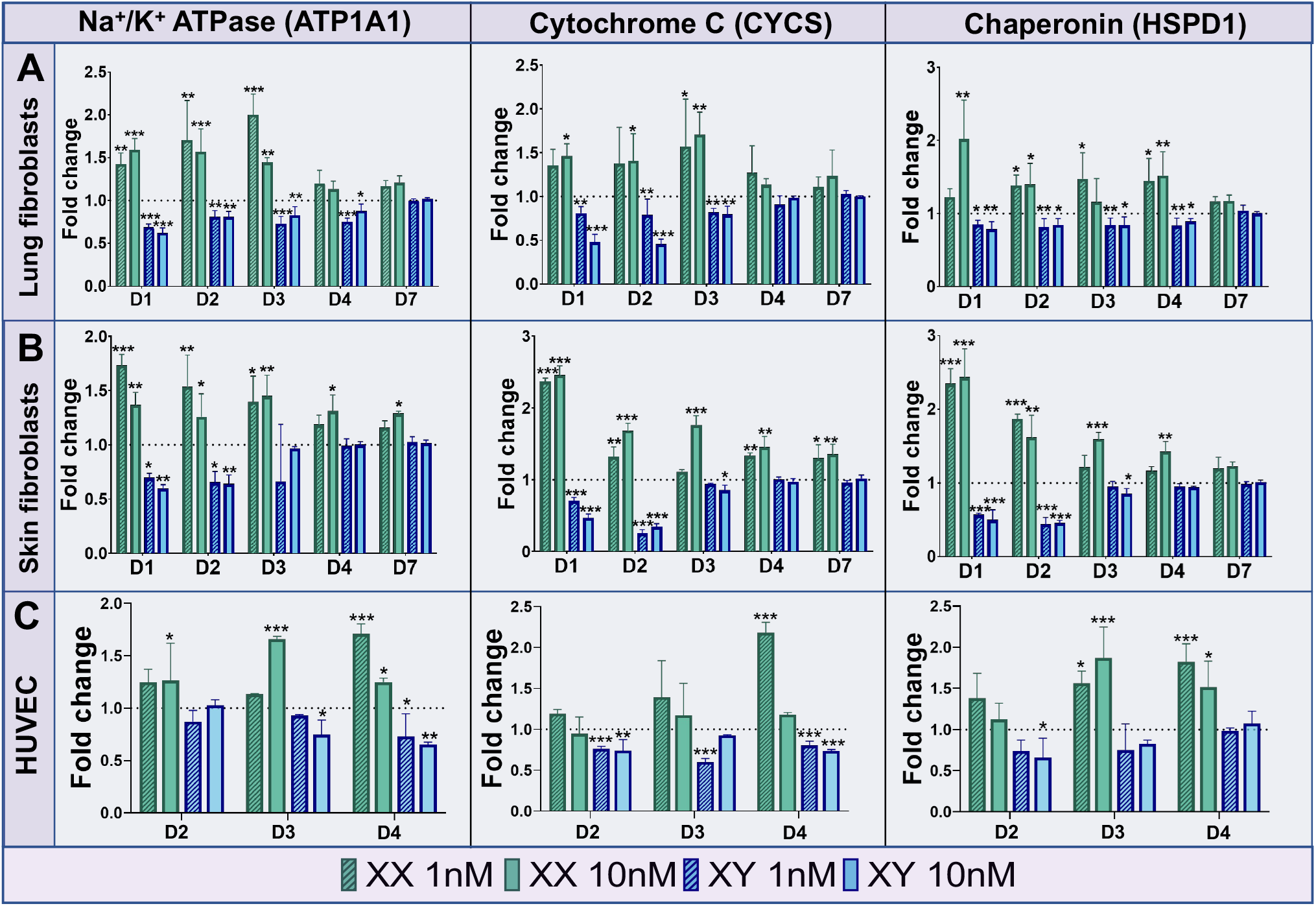
Sex-specific effects of E2 on endothelial cell and fibroblast expression of ATP1A1, CYCS, and HSPD1 mRNAs. **A:** PCR results for energetic and stress related genes in HLFs following exposure to varying E2 concentrations in XX (green) and XY (blue) cells over 7 days. Dotted line represents vehicle control group expression levels. XX cells significantly upregulated ATP1A1, CYCS, and HSPD1 following E2 exposure while XY cells underwent significant downregulation of the same genes. **B:** PCR results for the same genes in HDFs show a similar trend. **C:** HUVECs underwent the same changes in gene expression to a less profound extent over the course of 4 days. In all 3 cell types, expression levels returned back to near normal exposure by 7 days. Analyzed using the 2-□□Ct method. Compared against GAPDH as a housekeeping gene via one-way ANOVA (n=4 for HLFs, n=3 for HDFs and HUVECs). * = p < 0.05; ** = p < 0.01; *** = p < 0.001.

In line with the anti-proliferative effects measure in XY cells, E2 treatment induced transient downregulation of the same genes in XY HLF and XY HDF (**Figure 2B**). In HLF and HDF, CYCS and HSPD1 mRNA expression decreased to less than 50% of baseline after exposure to both 1nM and 10nM E2 for 24 hours, but it returned to near baseline levels by 72 hours and fully returned to baseline by 7 days (**Figure 2A, B**). E2 induced gene expression patterns were similarly inverse for XX and XY HUVEC, albeit with different kinetics than fibroblasts (**Figure 2C**). in XX HUVEC E2 increased the mRNA expression of ATP1A1 at 48, 72, and 96 hours, although only the 10nM E2 resulted in a significant upregulation at 24 hours. Expression of ATP1A1 mRNA steadily decreased in XY HUVEC at the same time points with significant decreases present in the 10nM E2 groups at 72 and 96 hours (**Figure 2C**). CYCS mRNA was upregulated in XX HUVEC at 24, 72, and 96 hours, with a significant upregulation seen with 1nM E2 at 96 hours. In XY HUVEC, both 1nM and 10nM E2 caused downregulation of CYCS mRNA until the 96-hour end point, and the same trends were noted in HSPD1 mRNA expression patterns. Both 1nM and 10nM E2 upregulated HSPD1 mRNA in XX HUVEC at 72 and 96 hours, and while only 10nM E2 induced a significant down regulation of HSPD1 mRNA, the trend continued with XY cells downregulating HSPD1 mRNA until 96 hours.

We used tetramethylrhodamine (TMRM), a convenient fluorescent indicator of mitochondrial membrane potential, to quantify the bioenergetic capacity of HLF and HUVEC in response to sex-matched and cross sex hormone stimulation. The TMRM dye accumulates in mitochondrial membranes at a rate proportional to the mitochondrial membrane potential, for which a 10-mV increase can double ATP production capacity (27, 33). We hypothesized that sex-specific hormone effects on cellular mitochondrial membrane potential will correlate with ATP1A1 and CYCS gene expression patterns induced at the same hormone concentrations. The TMRM signal intensity was similar for XX and XY HLF, indicating that mitochondrial membrane potential of XX and XY cells is similar in hormone-free culture, but qualitative differences in TMRM signal intensity were readily observable upon hormone stimulation for 48 hours (**Figure 3A, B**). E2 increased the TMRM signal in a dose-dependent fashion in XX HLF (**Figure 3A, C**) and HUVEC (**Figure 3B, D**). In contrast, E2 treatment decreased the TMRM signal in XY HLF (**Figure 3A, D**) and XY HUVEC (**Figure 3B, F**) in a dose-dependent fashion. Thus, E2 induced sex-specific alterations of mitochondrial membrane potential in HLF and HUVEC.

**Figure 3.**
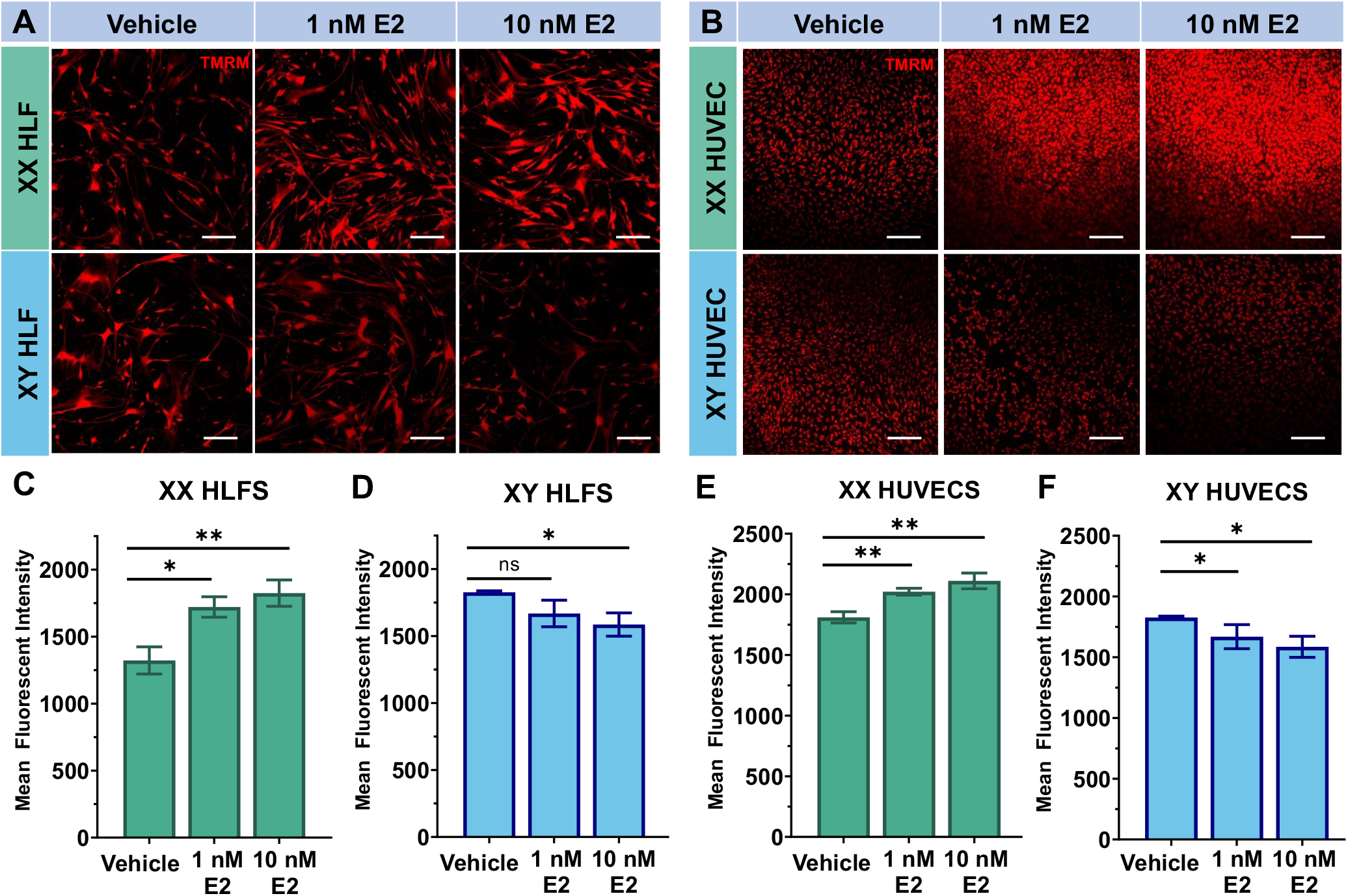
Sex-specific effects of E2 on endothelial cell and fibroblast mitochondrial membrane potential. **A:** Representative TMRM dye fluorescent micrographs of HLF monolayers comparing fluorescence activity between XX (top) and XY (bottom) exposed to 1nM E2, 10nM E2 or vehicle control for 48 hours. **B:** Representative TMRM dye fluorescent micrographs of HUVEC monolayers comparing fluorescence activity between XX (top) and XY (bottom) exposed to 1nM E2, 10nM E2 or vehicle control for 48 hours. Scale bar = 200 µm. **C-E:** Comparison of mean fluorescent intensity between groups shows a significant increase in fluorescence in XX HLFs following E2 exposure at both 1nM and 10nM (C) while XY HLFs underwent a significant decrease in fluorescence following E2 exposure (D). HUVECs followed similar trends with XX cells experiencing significant increase (E) and XY cells experiencing a significant decrease (F) Analyzed via unpaired, two-tailed t-tests based on n = 3 wells, 5 images per well.* = p < 0.05; ** = p < 0.01

### Dihydrotestosterone Induces Sex-specific Alterations of Cellular Bioenergetics in 2D Culture

We analyzed sex-specific responses of HLF and HUVEC to 1 and 10 nM DHT. We measured the same overall trends of sex-matched hormone and cross-sex hormone stimulation, but there were important differences in the magnitude and duration of the effects when compared to E2 treatment in the same cells. DHT treatment induced an approximately 2-fold upregulation of ATP1A1, CYCS, and HSPD1 mRNA in XY HLF that was sustained until the 7-day time point at both 1nM and 10nM DHT concentrations, in contrast to the return to baseline seen with sex-specific E2 stimulation in XX cells (**Figure 4A**). DHT treatment of XX HLF induced significant downregulations of these genes at both 1nM and 10nM that were also maintained until 7 days (**Figure 4A**, green bars), indicating more persistent inhibitory cross-sex effects of DHT compared to E2 in the same cell culture assays. We measured similar gene expression patterns in DHT treated HUVEC at both 1nM and 10nM DHT, except for CYCS, which returned to baseline in XY HUVEC by 7 days and HSPD1 which returned to baseline in 10nM DHT by 7 days (**Figure 4B**). Taken together, these analyses demonstrate that E2 and DHT exert similar sex-specific effects in defined culture medium and highlight the need for more rigorous development of sex-specific culture systems and methods.

**Figure 4.**
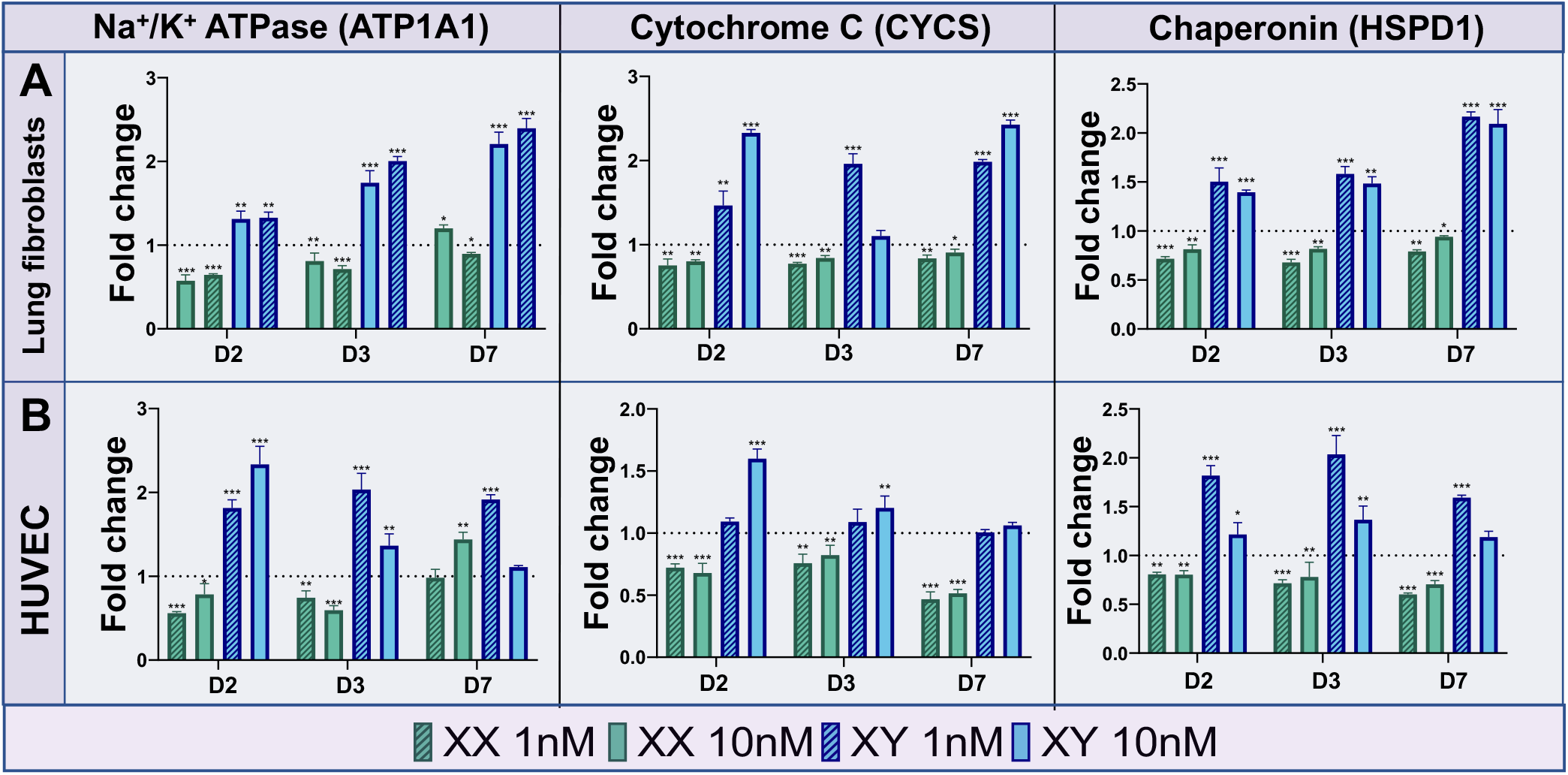
Sex-specific effects of DHT on endothelial cell and fibroblast expression of ATP1A1, CYCS, and HSPD1 mRNAs. **A:** PCR results for energetic and stress related genes in HLFs following exposure to varying DHT concentrations in XX (green) and XY (blue) cells over 7 days. Dotted line represents vehicle control group expression levels. XX cells significantly downregulated ATP1A1, CYCS, and HSPD1 following DHT exposure while XY cells underwent significant upregulation. **B:** PCR results for the same genes in HUVECs show a similar trend. Analyzed using the 2-□□Ct method. Compared against GAPDH as a housekeeping gene via one-way ANOVA (n=3). * = p < 0.05; ** = p < 0.01; *** = p < 0.001

Consistent with gene expression trends, DHT qualitatively increased the TMRM signal in XY HLF, with a converse dose-dependent decrease in XX cells (**Figure 5A**). We observed similar qualitative patterns of TMRM signal intensity for DHT treated XX and XY HUVEC (**Figure 5B**). Quantifying the mean fluorescence intensity revealed that DHT treatment significantly increased the TMRM signal in XY HLF at 1 (p<0.05) and 10 nM (p<0.05) (**Figure 5D**). Conversely, the TMRM signal intensity decreased significantly when XX HLF were treated with 1 (p<0.05) and 10 nM DHT (p<0.01) (**Figure 5C**). DHT treatment similarly decreased the mitochondrial membrane potential of XX HUVEC, but the magnitude of the effect was not as pronounced when compared to XX HLF with only 10nM DHT inducing a significant decrease (p<0.05) (**Figure 5E**). 1 nM DHT significantly increased the TMRM signal intensity in XY HLF (p<0.05), but 10 nM DHT significantly decreased the signal intensity (p<0.05) (**Figure 5F**).

**Figure 5.**
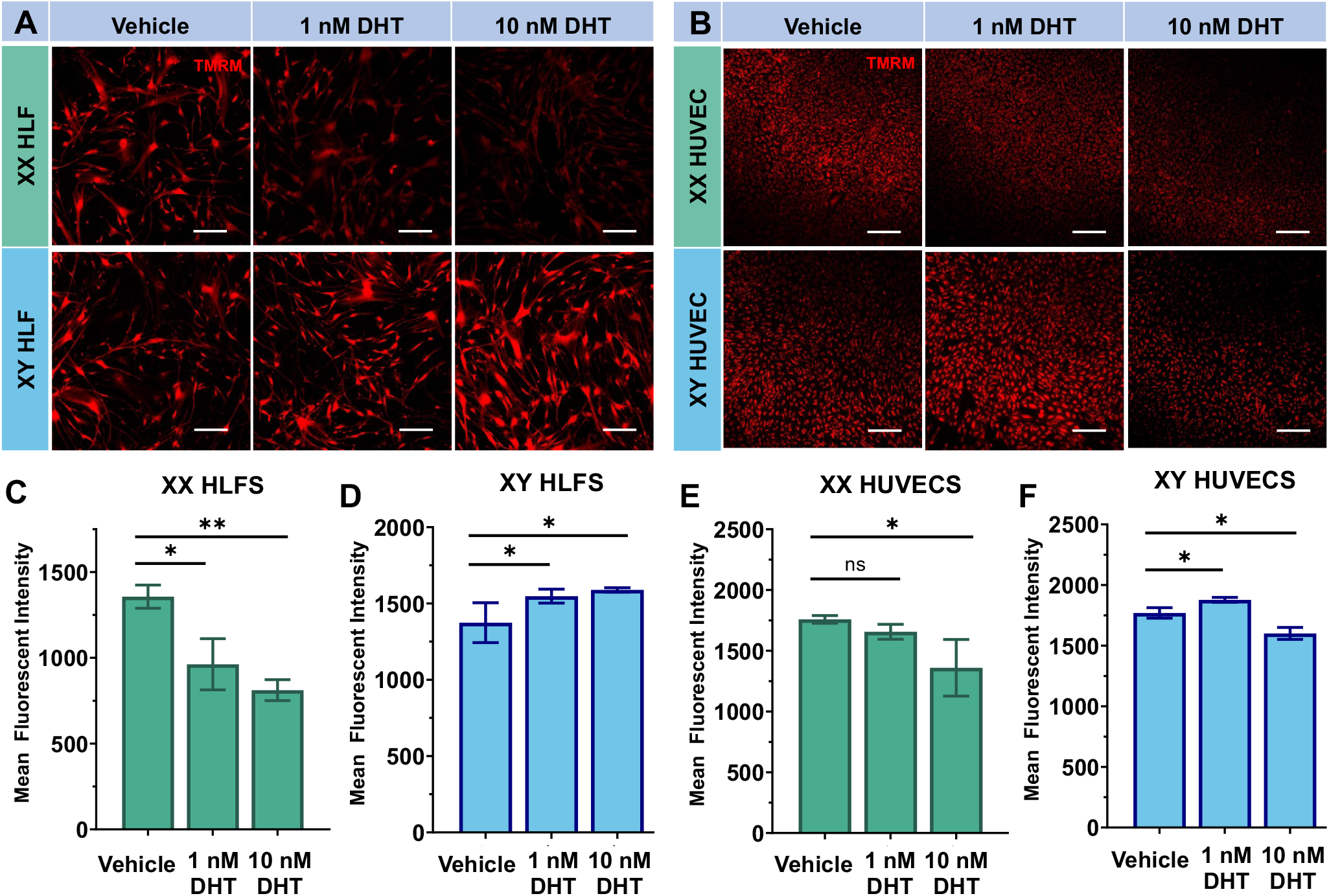
Sex-specific effects of DHT on endothelial cell and fibroblast mitochondrial membrane potential. **A:** Representative TMRM dye fluorescent micrographs of HLF monolayers comparing fluorescence activity between XX (top) and XY (bottom) exposed to 1nM DHT, 10nM DHT or vehicle control for 48 hours. **B:** Representative TMRM dye fluorescent micrographs of HUVEC monolayers comparing fluorescence activity between XX (top) and XY (bottom) exposed to 1nM E2, 10nM E2 or vehicle control for 48 hours. Scale bar = 200 µm. **C-E:** Comparison of mean fluorescent intensity between groups shows a significant decrease in fluorescence in XX HLFs following DHT exposure at both 1nM and 10nM (C) while XY HLFs underwent a significant increase in fluorescence following DHT exposure (D). HUVECs followed similar trends with XX cells demonstrating a significant decrease at 10nM DHT (E) and XY cells experiencing a significant increase in TMRM intensity at both 1nM and 10nM DHT (F) Analyzed via unpaired, two-tailed t-tests based on n = 3 wells, 5 images per well.* = p < 0.05; ** = p < 0.01

### Sex-Specific Effects of E2 and DHT on Vasculogenesis in XX and XY Tissues

We next examined if the sex-specific effects of hormone stimulation on cell proliferation and bioenergetics would translate to 3D bulk tissue vasculogenesis. We designed a device that enables culture of 3D disc-shaped tissues with microengineered surface anchorage to maintain a fixed volume and geometry (**Figure 7A**). The devices are composed of polydimethylsiloxane (PDMS), a common material used in organ chip applications (34). Of particular importance, PDMS is hydrophobic and therefore absorbs small hydrophobic molecules such as E2 and DHT (24). We employed surface coating methods to render PDMS surfaces hydrophilic and therefore resistant to hormone absorption (**Figure 6**) (25). Nile Red absorption studies were completed on plain PDMS and PDMS PEG copolymers cast on both clear coated and noncoated molds (**Figure 6A-D**). We quantified the average gray intensity across the device in FiJi (**Figure 6E, F**), and observed a higher fluorescent signal in devices made on clear vs unclear coated molds (p<0.001). The presence of the PEG copolymer further decreased the fluorescence signal of the Nile Red (p<0.001) (**Figure 6F**). Hence, the low level of Nile Red absorption in the copolymer devices formed on clear coated molds suggested that this preparation is sufficient to avoid absorption of E2 and DHT by the microarrays, and this composition was used for all further vasculogenesis studies.

**Figure 6.**
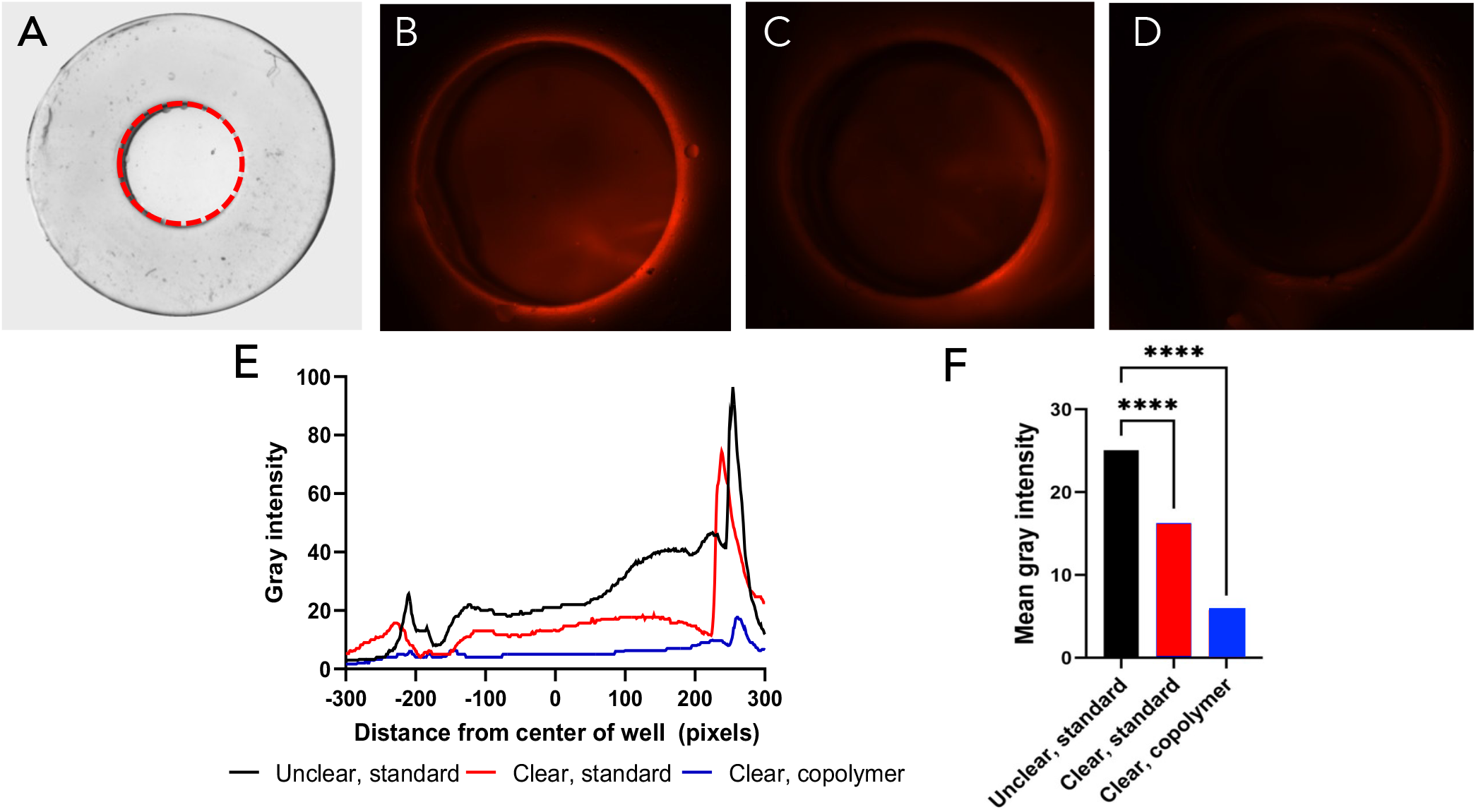
PDMS modification to mitigate bulk hormone absorption. **A:** Brightfield image of sample PDMS disc. Area of interest is the well in the center outlined with red dashed line. **B-D:** Fluorescence microscopy images of area of interest of PDMS discs after exposure to 1 ug/ml Nile Red for 1 hour. Standard PDMS molded on an unclear coated mold (B), standard PDMS molded on a clear coated mold (C), and PDMS+PEG Copolymer molded on a clear coated mold (D) were tested. **E:** Linear fluorescent intensity across PDMS discs. Lines represent average value of 3 discs. **F:** Comparison of mean gray intensity shows a significantly higher fluorescent signal corresponding to significantly increased absorption in standard, unclear PDMS compared to standard, clear PDMS. PDMS+PEG copolymer showed significantly decreased absorption.

**Figure 7.**
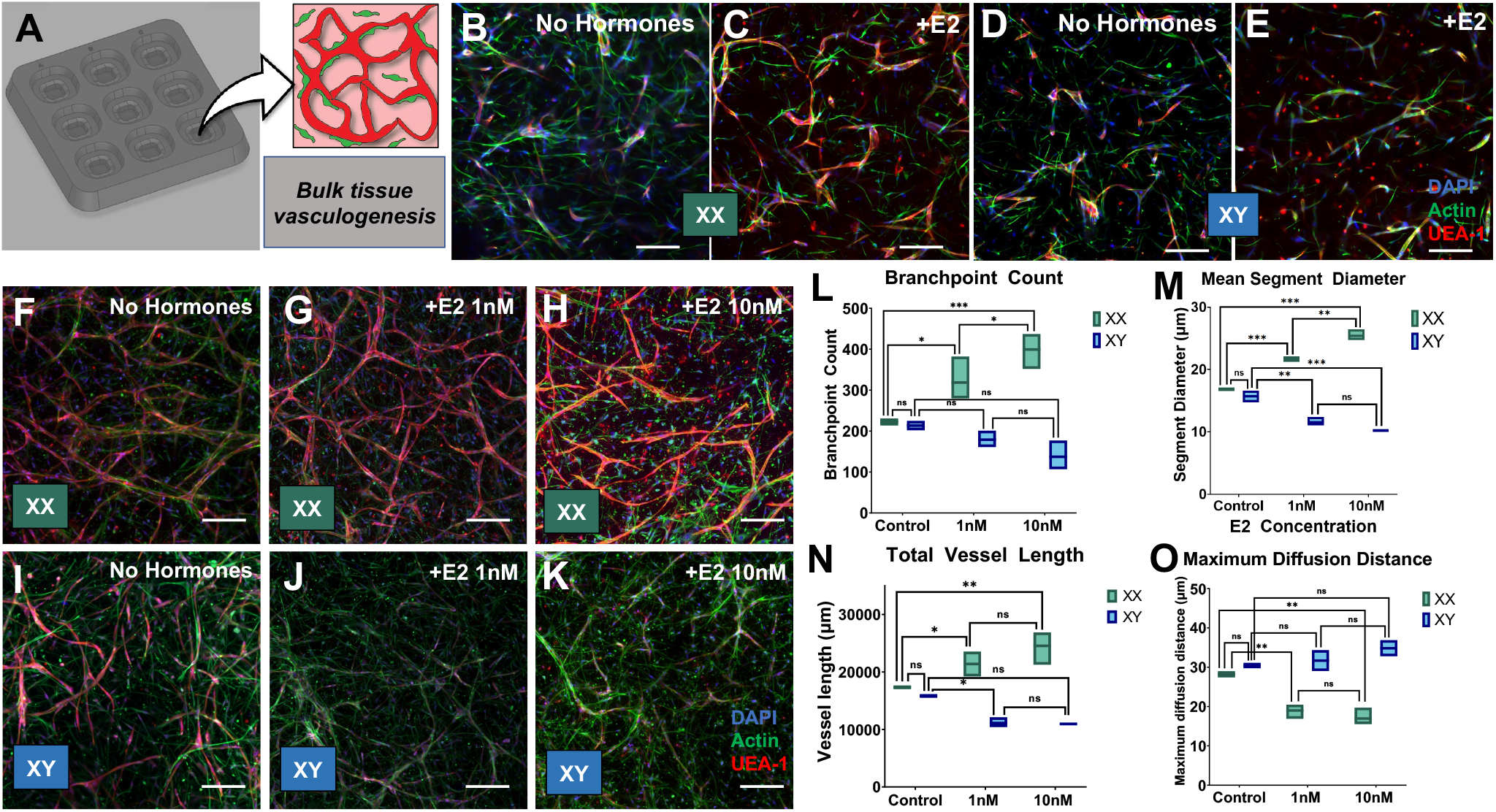
Sex-specific effects of E2 on bulk tissue vasculogenesis. **A:** Solidworks renders of PDMS microarray device in which vasculogenic gels are loaded and allowed to undergo bulk tissue vasculogenesis. **B-E:** Representative micrographs of XX tissues and XY tissues cultured for 2 days in hormone stripped media (**B and D** respectively) demonstrate similar levels of vascularization. Gels cultured with 10nM E2 for 3 days show differences between XX and XY gels. XX gels exposed to E2 exhibit increased UAE-1 levels and more complex vessel formation (**C**) while XY gels exposed to E2 show rounded endothelial cells stained in UAE-1 and significantly lower vessel formation (**E**). **F-K**: XX and XY devices grown in 1nM or 10nM E2 for 5 days following the 2-day starvation period show similar differences in network complexity. Some vessel formation is seen in XY gels at day 7 (**J and K**), but XX gels exposed to 1nM and 10nM E2 show more organized UAE-1 in complex vascular networks (**G and H** respectively). Scale bars = 200 µm. L-O: Morphometric outputs generated by MATLAB analysis show that XX networks have significantly higher branchpoint counts (**L**), mean segment diameter (**M**), and total vessel length (**N**) along with lowered mean diffusion distance (**O**). The opposite trends are seen in XY gels. Compared via two-way ANOVA. (n = 4) * = p < 0.05; ** = p < 0.01; *** = p < 0.001

A bulk tissue vasculogenesis assay for these studies was created by forming 3D tissues comprised of HLF and HUVEC embedded in a blended collagen type I and fibrin hydrogel. We treated XX and XY tissues with 1 nM or 10 nM E2 or 1nM or 10nM DHT and compared metrics of vascular network formation with baseline culture conditions (stripped serum, no hormones). Vasculogenesis was assessed by co-staining with the endothelial-specific lectin *Ulex Europeas* Agglutinin-1 (UEA-1) and phalloidins to label F-actin in all cells (**Figure 7B, D**). Control tissues were treated with vehicle at the same concentration as hormone-stimulated cultures.

We first observed that 10 nM E2 enhanced vasculogenesis relative to baseline by 48 hours (**Figure 7C compared to B**). In contrast, and consistent with anti-proliferative effects in XY HUVEC (**Figure 1**), E2 potently abrogated vasculogenesis in XY tissues and many endothelial cells appeared rounded, suggesting that E2 negatively regulates endothelial cell adhesion and spreading in XY cells (**Figure 7E compared with D**). We next treated XX and XY tissues with 1 nM or 10 nM E2 for 5 days and used total vessel length, mean segment diameter, total branchpoint count, and maximum diffusion distance as metrics to quantify vasculogenesis. There were no significant differences in the metrics mentioned above between XX and XY tissues in baseline cultures (no hormones, stripped serum), demonstrating that there are no intrinsic differences in the vasculogenic capacity of XX and XY tissues in this 3D culture configuration (**Figure 7L-O**). E2 increased branchpoint count, mean segment diameter, and vessel length, and decreased the maximum diffusion distance in XX tissues (**Figure 7F-H and L-O**). This effect was seen at both 1nM and 10nM E2 with 10nM E2 inducing a more significant influence. Conversely, E2 decreased vessel length, branchpoint count, and mean segment diameter, and increased the maximum diffusion distance in XY tissues at both 1 and 10nM E2 (**Figure 7I-K and L-O**). Thus, E2 stimulates a robust vascular network formation in XX tissues and potently inhibits vasculogenic capacity in XY tissues. These data demonstrate that sex-specific effects of E2 on proliferation and bioenergetic capacity of primary endothelial cells and fibroblasts described above correlate with the stimulatory and inhibitory tissue-scale effects of E2 on vasculogenesis.

In contrast to E2 exposures, DHT did not induce significant changes in any of these metrics in either XX or XY tissues (**Figure 8A-J**). Instead, we measured a significant increase in rounded, non-participating endothelial cells in XX tissues exposed DHT compared to XY tissues (**Figure 9**). Interestingly, DHT also increased the number of non-participating endothelial cells in XY tissues relative to baseline (**Figure 9**). These data suggest that DHT at 1 and 10 nM may inhibit cell-ECM adhesions or intercellular adhesions in endothelial cells. Thus, E2 and DHT exert divergent sex-specific effects on bulk tissue vasculogenesis. Collectively, our findings establish a clear methodology for using E2 to enhance vascularization of XX tissues, whereas the use of DHT to optimize vascularization of XY tissues requires further investigation.

**Figure 8.**
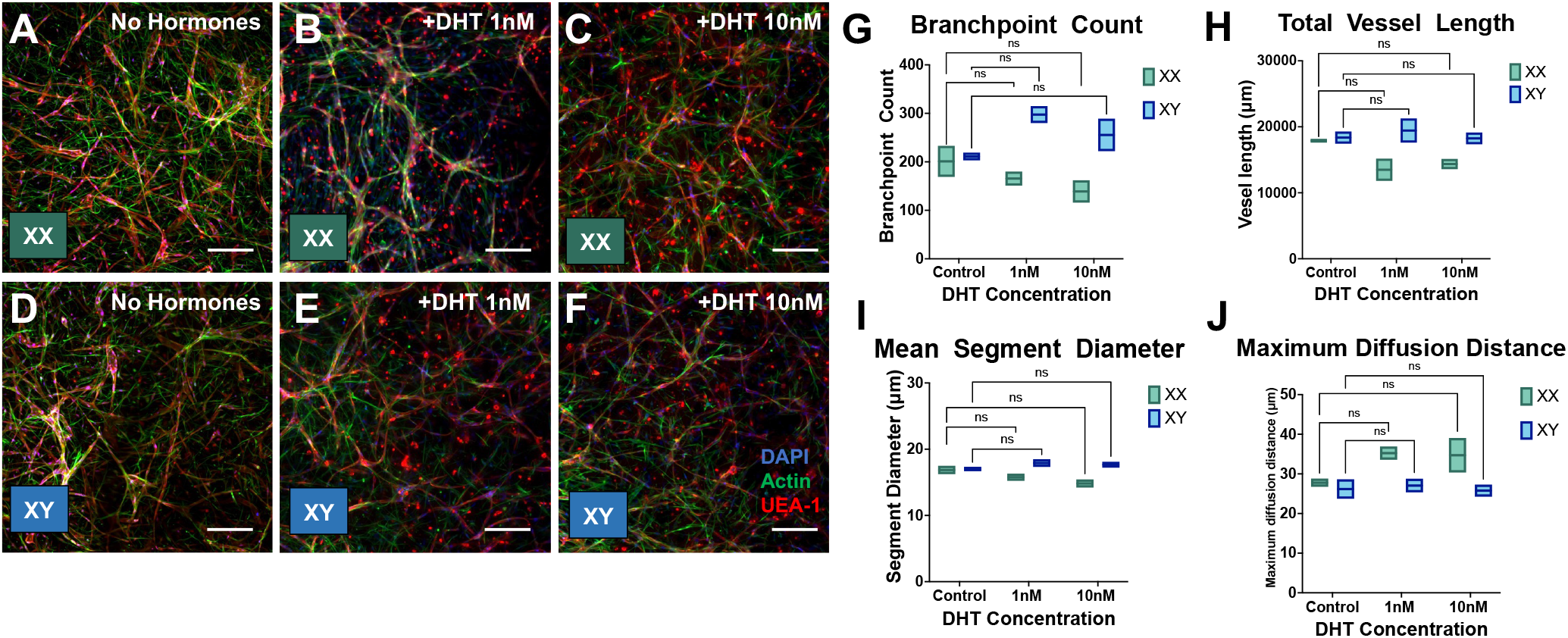
Sex-specific effects of DHT on bulk tissue vasculogenesis. **A-F:** Representative micrographs of XX (top) and XY (bottom) tissues cultured for 2 days in hormone stripped media followed by 5 days of 1nM or 10nM DHT exposure. Control groups show similar network formation. Gels grown in 1nM and 10nM DHT displayed similar vessels but with more UEA-1 positive cells that have a rounded morphology and are not in the vascular network. Scale bars = 200 µm. **G-H:** Morphometric outputs generated by MATLAB show that there was not significant difference between XX and XY gels grown in DHT versus control samples. Compared via two-way ANOVA (n=3). K-L: As evidenced by the increase in rounded epithelial cells stained for UEA-1, DHT caused a significant increase in non-participating cell counts in both XX and XY tissues while E2 only caused an increase in non-participating cells in XY tissues. Compared via two-way ANOVA. (n = 4) * = p < 0.05; ** = p < 0.01; *** = p < 0.001

**Figure 9.**
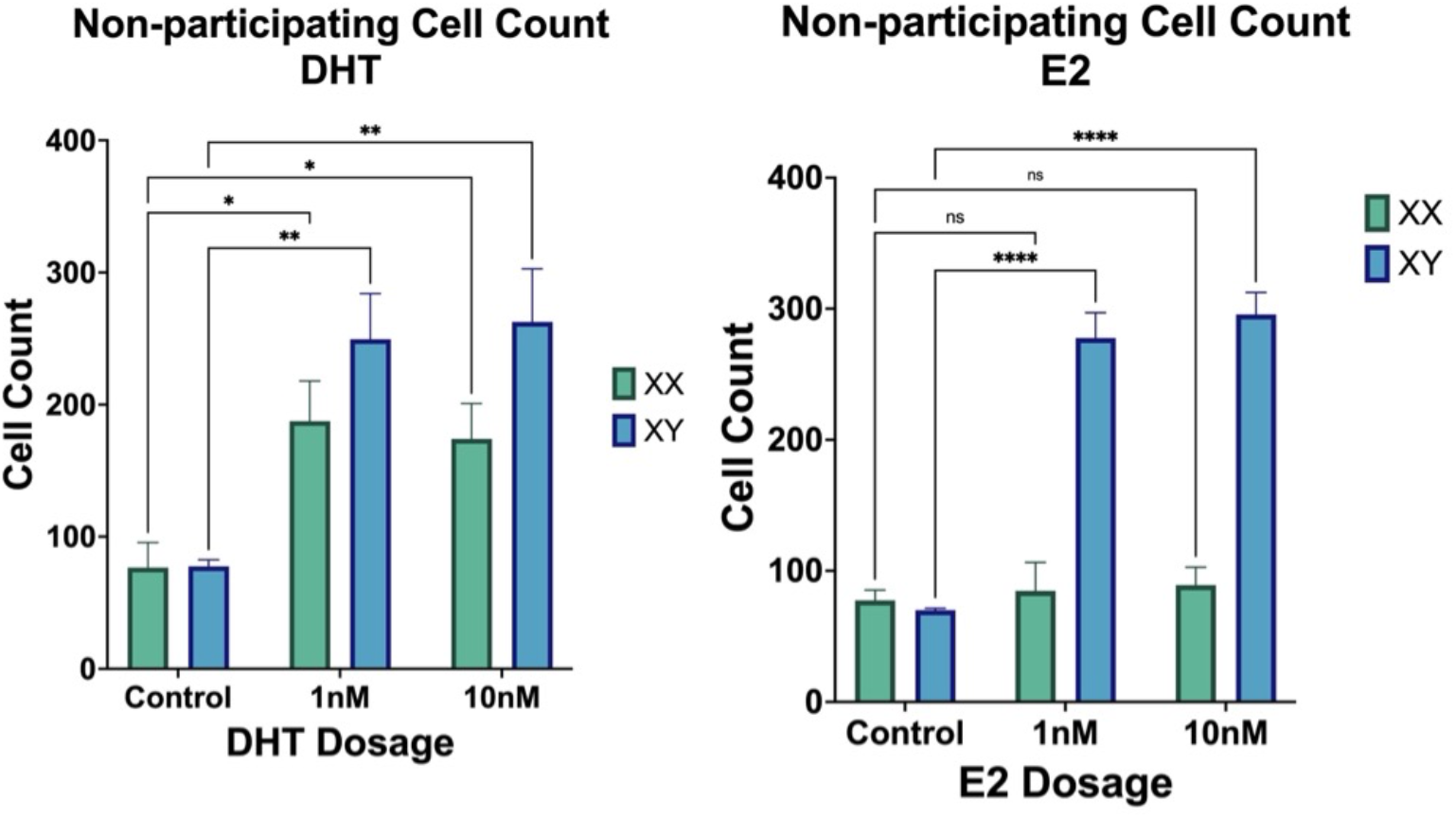
Numbers of non-participating endothelial cells in XX and XY tissues treated with DHT or E2 in the vasculogenesis assay. DHT increases non-participating endothelial cell numbers in both XX and XY tissues, whilst E2 has no effect.

## Discussion

### Designing male and female cell culture environments

The development of physiologically relevant tissue engineering technologies and microphysiological systems requires careful consideration of individual donor differences during model development. Biological sex is a critical variable that contributes to the heterogeneity of cellular responses in engineered tissue systems along with age, race, lifestyle, and comorbidities. The concept of heterogeneous responses based on intrinsic cellular properties is commonly applied when creating disease models using cell cultures derived from patients with the disease compared with cells from control subjects without the disease, yet these studies typically fail to stratify results based on biological sex (35, 36). This lack of consideration of SABV in these areas of biotechnology development is surprising as sex is a genetic modifier of biology and disease (3, 37, 38).

We established that sex-specific hormone stimulation, i.e., E2 with XX cells and DHT with XY cells, enhances cellular bioenergetics and increases rates of proliferation of primary human fibroblasts and endothelial cells. We believe this is the first study to compare E2 and DHT effects on proliferation and bioenergetics in female and male human primary cells under identical experimental conditions. Our findings have important implications for the development of sex-specific biotechnology processes. For example, expansion of primary human cells is a key challenge in all aspects of tissue engineering from the creation of in vitro models to and cell therapies and macroscopic organ engineering (39, 40). The importance of sex hormones in culture medium design has been discussed as an area requiring attention, but there are no widely adopted protocols for cell and tissue culture based on the biological sex of the cells being cultured (41). Fetal bovine serum (FBS) is a common component of cell culture media formulations used to drive proliferation of cultured cells. The nature of FBS manufacturing and inter-lot variability of FBS results in an unpredictable mixture of bovine estrogens and testosterone, with both hormones present in physiologically relevant amounts (42–44). For instance, in FBS from 4 different suppliers, E2 concentrations ranged from 58 to 232 pg/ml (45). Furthermore, serum-free medium formulations do not account for the loss of sex hormones with the removal of serum. Artificial biological aging and senescence are major challenges that come with repeated cycles of cell division, depending on the mitogenic stimulus (46, 47). Sex hormones may provide a means of driving proliferation in a manner that preserves biological age, as has been demonstrated for human hematopoietic stem cells (48).

### Bioenergetics drive sex-specific hormone effects in expansion culture

The ability of cells to produce ATP dictates proliferation in physiological and pathological contexts (49, 50). Here, we provide evidence that E2-induced proliferation of XX cells and DHT-induced proliferation of XY cells is driven at least partially by increased bioenergetic capacity (**Figure 2, 3, 4, and 5**). Similarly, our studies provide evidence that anti-proliferative cross-sex hormone effects (**Figure 1**) are driven by a decreased bioenergetic capacity. Previous research has established that sex hormones modulate bioenergetic capacity in various contexts, but there are no studies that compare sex-matched and cross sex hormone stimulation of XX and XY donor cells under the same experimental conditions. E2 is a positive regulator of cellular bioenergetics in multiple female organ systems via regulation of glucose metabolism and mitochondrial production of ATP. For instance, E2 has positive effects on all stages of ATP production in the brain (51, 52), and declining estrogen levels result in overall reduced brain bioenergetics (53). Similarly, DHT increases cell proliferation, protein synthesis, and energy metabolism in male tissues and cells including skeletal muscle (54).

### Cell type specificity of responses to sex hormone stimulation

We found that DHT effects more pronounced in HLF, whereas E2 effects were more pronounced in HUVEC (**Figure 1-5**). Notably, the inhibitory effect of E2 on XY tissue vasculogenesis was seen more prominently in the large numbers of rounded or poorly spread endothelial cells, whilst HLF in the same tissues exhibited a typical elongated morphology and distribution (**Figure 7**). These observations imply that cell lineage is a critical factor that impacts the response to sex hormone stimulation in culture. Future development of sex-specific culture systems, for example culture medium formulations, can be accelerated and improved by a deeper understanding of the interplay between biological sex and organ-specific physiology.

Hormone responses of cells such as endothelial cells and fibroblasts will vary depending on the organ of origin (55). The similar response of HLF and HDF upon E2 treatment suggests a potentially conserved mechanism by which estrogen regulates interstitial tissue homeostasis throughout the body in biological females (**Figure 2**). On the other hand, there are reports of organ-specific fibroblast responses to E2 stimulation (23). While some aspects of sex hormone regulation of fibroblast physiology may be conserved, it is possible that organ-specific effects of sex hormones may emerge in the context of pathology or tissue injury. Future studies should test the response to E2 and DHT in fibroblasts and endothelial cells derived from a larger number of organs to investigate organ-specificity of hormone responses in greater depth. More broadly, our study highlights that caution should be taken before extrapolating results from a given cell type to predict responses of a different lineage in culture.

### Cross sex hormone stimulation: toward models of gender-affirming hormone therapy

Our cross-sex hormone stimulation data warrants further investigation in the context of cross-sex hormone therapy (CSHT), also known as gender-affirming hormone therapy. To the best of our knowledge, there are no research reports about developing cell culture models of CSHT. PubMed searches for the term ‘transgender hormone therapy cell culture’ and related variations thereof return only a few citations with overlapping keywords, none of which involve in vitro experimentation aimed at understanding cross sex hormone effects in human cells. Information from cell culture systems could validate established clinical trends such as increased cardiovascular disease in transgender individuals (56–59), while potentially improving our understanding of controversial clinical trends. For example, one study reported a statistical increase in cardiovascular disease in transgender females on estrogen therapy, whereas no change in incidence was reported for transgender males on testosterone therapy (60). Whilst our simplified hormone culture experiments do not comprehensively mimic the physiology of individuals on CSHT or emergent clinical trends, our findings that E2 exerts inhibitory effects on XY cell proliferation, bioenergetic capacity, and bulk tissue vasculogenesis clearly demonstrates potentially pathological effects of cross sex hormone stimulation (**Figure 1, 2, 3, and 7**). Thus, human cell-based engineered systems can potentially be used to investigate the mechanisms underlying clinical observations such as the increased risk of cardiovascular pathology in transgender women.

In addition, our finding that DHT suppresses cellular bioenergetics in XX cells corroborates prior preclinical research. For example, exogenous DHT decreased placental mitochondrial content and ATP generation capacity in a rat model (61). Our qualitative observation that cross sex hormone stimulation inhibits actin cytoskeleton assembly in HLF warrants further investigation. Our data clearly demonstrates that cross sex hormone stimulation decreases bioenergetic capacity in short term culture (**Figure 3**, **5**), but there may be direct signaling effects that negatively regulate cytoskeletal dynamics independently of energetic state. In support of altered cell mechanics upon cross sex hormone stimulation, we observed that XY endothelial cells in tissues were rounded after 48 hours of E2 treatment, which could potentially render these cells susceptible to cell death by anoikis. Furthermore, the trend of decreased chaperonin expression with cross sex hormone stimulation in 2D cultures of all cell types tested suggests the possibility of increased sensitivity to endoplasmic reticulum stress (62). Future studies should investigate whether the decrease in cellular bioenergetic capacity in cross sex hormone-treated human cells translates to an increased sensitivity to proapoptotic stimuli.

### Future directions for engineering biologically sex-matched tissues

The results of our vasculogenesis study highlight the challenges that lie ahead for the comprehensive development of sex-specific tissue and organ models. Whilst the enhanced vasculogenic response of XX tissues to E2 at the applied concentrations sets a clear path for using estrogens to enhance female vascular tissue engineering (Figure 7), the inhibition of vasculogenesis in XY tissues stimulated with DHT (Figure 8) suggests that using androgens to enhance male vascular tissue engineering is not as straightforward. It is possible that lower concentrations of DHT are more physiological, i.e. in the 0.1 nM range; however, much investigation is needed to address this issue. There are many limitations of the current study. Most notably, the female and male hormone environments need to be more refined to accurately mimic physiology. Progesterone (P4) effects alone and in combination with E2 need to be elucidated. E2 should be added to male culture medium, and DHT should be added to the female medium, both at appropriately low levels to mimic in vivo concentration ranges. Furthermore, the epigenetic landscape of hormone stimulated cells in culture should be characterized and compared with freshly isolated in vivo counterpart cells to optimize hormone concentrations in sex-specific culture systems for the preservation of physiological characteristics. Upon establishment of more physiomimetic hormone formulations for male and female culture media, the effects of sex-matched and cross-sex hormone stimulation need to be investigated in a broader range of cell types, for example epithelial tissues of endodermal origin, and neuronal tissues of ectodermal origin. Lastly, continued development of sex-specific models will enable investigation of hormone status effects in microphysiological models of diseases such as cancer and diabetes.

## Declarations

The research reported in this paper was funded by Mark J. Mondrinos’ faculty startup package provided by Tulane University and pilot investigator funds from the Tulane Center for Sex-based Biology and Medicine (TCESBM). FMJ was supported by National Institutes of Health grant DK074970, a U.S. Department of Veterans Affairs Merit Award BX003725 and the TCESBM. The authors have no conflicts of interest to report. All CAD design files, and the code used for image analyses are made available in the Supplementary Information for this article. All data and more detailed explanations of laboratory procedures will be made available upon reasonable request to the corresponding author.

